# The correlation between Rate of Force Development Maximal Strength and Electromyography Variables of Basketball Athletes

**DOI:** 10.1101/2024.01.31.578254

**Authors:** Hiury Caio Pinheiro Brandão, Marcelino Monteiro de Andrade, Jake Carvalho do Carmo

## Abstract

The rate of force development (RFD), is seen as a determining characteristic in fast actions present in basketball. However, we observed different relationships between RFD and maximum strength, as well as different relationships between RFD and neuromuscular variables according to the evaluated population. The aim of the present study is to evaluate the degree of determination of maximum strength (Tmax) and neuromuscular recruitment variables (RMS), Absolute Energy (AE) and the motor units firing frequencies (MPF) in rate of force development (RFD) for basketball athletes. Nine basketball athletes from the same team (mean ± SD; age: 20.8 ± 2.08 years; body mass: 84.33 ± 8.80kg; height: 1.86 ± 0.095 meters; practice time: 11.67 ± 1.65 years) were evaluated through maximum isometric contraction with highest value of maximum force among 3 attempts. The RFD were evaluated and correlated with the RMS and AE values and the MPF values of the electromyographic signal at instants 0-50; 50-100, 100-150 and 150-200 milliseconds. The results show a reduction in RFD and MPF over the evaluated time windows and also a correlation between MPF and TDF in the 0-50ms time window (R^2^ 0.67 *p*<0.05). The results show no relationship between RFD and RMS and AE, in addition to these variables not showing significant reductions in the evaluated time windows. The levels of RFD show to be more related to the firing frequency of the motor units than the maximum force and the level of recruitment of the motor units.

## Introduction

Muscular strength is one of the most important physical qualities when it comes to sports, as it is directly related to performance of the sport itself. In this sense, several authors have corroborated the idea that the employment of force differs on each sport. It is possible to identify muscular strength applied at maximum levels – known as “maximum strength” – and those with fast development – known as “rate of force development” [1-3].

The relationship between maximum strength and the rate of force development (RFD) may present different behaviors according to the population evaluated. While previous studies have shown moderate and strong significant correlations between these forces characteristics among the sedentary population, such relationship has not been identified in the evaluation of football athletes as in national-level cycling athletes. These studies suggest that RFD may be an independent variable to maximum strength among the athletes’ population [4,5,6].

As an example, while playing basketball, the athlete’s ability to produce maximum strength is directly linked to the functional actions of the sport. There is a correlation between the performance of maximum strength and the performance in specific tasks such as jumps, sprints and agility tests, as well as a correlation between the RFD that is produced during a vertical jump [7,8].

Despite the correlation between the ability to produce maximum strength and the level of performance of motor tasks that are specific to basketball, it is possible to notice a short period of time when the production and application of strength in the movements of the sport occurs. For instance, when carrying out sprints and jumps at considerable heights, the time of application of strength to the ground have a specific duration, respectively, in the range of 80-100ms and 170-180ms. These times are not enough to reach the maximum strength that is possible to be produced by an athlete, as the production of maximum strength and strength development are characterized in sports movements as time-dependent [9,10].

Strength production in basketball is characterized by the Rate of Force Development (RFD) curve. This quantity consists of the variation of force by the variation of time in short windows [11].

RFD can be divided into two phases: initial and late. The initial force development occurs in the first 100ms of the movement whereas the late force development shall occur after that period. These phases seem to be driven by different factors: while the initial moments are more connected to neural activation factors, the late moments are more related to morphological and mechanical aspects [12].

The recruitment of motor units is initially presented as a determinant of the initial RFD. It is possible to mention two more determining factors in order to improve the explosive strength of people who perform strength training. Initially, we shall observe the amplitude of the electromyographic signal (RMS) in initial time windows and then, the speed of recruitment of the units. Such aspects are indicated by the percentage of the RMS value in each window compared to the maximum RMS. [13]

However, previous investigations in physically active participants have identified that not only does the speed of recruitment of motor units is determinant for the development of initial RFD, but there is also a large contribution of the maximum rate of firing in these motor units. This rate was calculated by the median power frequency (MPF) of each window [14].

It is known that maximum strength is trained in a hierarchical way before the other characteristics of muscle strength, which demands a long time of preparation for an athlete. It is also noted that maximum strength in basketball, despite not having the necessary time to be reached during its movements, does have a correlation with the performance of the fundamental movements of the sport (sprint, jumps and speed of change of direction) [15].

Thus, it is not yet possible to identify whether the maximum strength is determinant for the RFD in basketball athletes. Additionally, it is not yet possible to establish which neural factor is determinant for the optimization of the rate of force development. Therefore, the aim of the present study is to evaluate the degree of determination of maximum strength and neuromuscular recruitment variables (RMS) and the frequency of firing of motor units (MPF) in rate of force development (RFD) for basketball athletes.

## Materials and methods

### Participants

Nine participants that are basketball athletes volunteered for this study (mean ± SD; age: 20.8 ± 2.08 years; body mass: 84.33 ± 8.80kg; height: 1.86 ± 0.095 meters; practice time: 11.67 ± 1.65 years). Everyone trained on the same team. The free and informed consent form was signed by all participants and the study was conducted in accordance with the Declaration of Helsinki. The approval for this project was granted by the Ethics and Research Committee at the Faculty of Health Sciences of the University of Brasília – UnB, under number 2,197,665.

### Experimental Procedure

Participants were initially prepared for data collection by identifying their dominant lower limb through the test question “were you to kick a ball at a target, which leg would you choose?” [16]. The participant was then directed to the placement of the surface electrode. In order to evaluate the electromyographic variables, the vastus lateralis muscle was chosen. Hair trichotomy, skin abrasion and cleaning with 70% alcohol and cotton were performed. The demarcations of the muscular belly zone and the placement of the electrodes were all performed under the guidelines of SENIAM [17].

Next, the participants were directed to perform the maximum voluntary isometric contraction (MVIC) test on the isokinetic dynamometer (BIODEX® Model System 4). Within the exercise protocol, a warm-up consisting of (*i*) 2 sets x 10 repetitions with an angular velocity of 120 º/s for the concentric phase of knee extension and (*ii*) an angular velocity of 300 º/s for the knee flexion phase was performed. Along with the warm-up, a series of isometric sub-maximal contraction at 60º of 3 seconds were performed for familiarization, even though the volunteers were already acquainted with the test, as it was periodically performed within their team, and the equipment. After each series of warm-ups, an interval of 30 seconds was performed.

After the warm-up, an interval of 5 minutes before the MVIC test was adopted. For MVIC collection, the knee angulation was positioned at 60º with the knee fully extended as point 0. This angulation was adopted because it is considered to be the degree of highest torque production, besides being considered the angle with the greatest muscle activation for the capture of electromyography signals [18,19].

To perform the data acquisition in the MVIC, the participant was asked to perform the knee extension movement as fast and as strong as possible, sustaining this force for a period of 3 seconds, when hearing the sound signal emitted by the isokinetic dynamometer to start until hearing the sound to finish. The MVIC protocol was performed with the ballistic movement and with the limb relaxed in the initial evaluation position, in 3 attempts, with an interval of 3 minutes between each. The highest performance was then considered for analysis.

## Measurement and data analysis

### Electromyography

The acquisition of electromyographic signals was performed using the *Delsys Bagnoli-2 EMG System* electromyograph and its electrodes. The device was set with amplification for electromyography signals of 1000 times, common rejection mode of 110 dB. The digital analog plate was programmed to receive the data with a sampling frequency of 2000 Hz.

The processing of electromyography data was carried out in the *Matlab2018*® environment. The values of the signal energy were calculated using the RMS and the Absolute Energy of the signal (AE), which was calculated through the sum of the absolute values of the electromyographic signal. The calculation was performed in the following windows: 0-50; 50-100; 100-150 and 150-200ms. Finally, in order to analyze the frequency data of the shots of the motor units, the calculation of the MPF was performed in the same windows for the analysis of the RMS.

### Torque and Rate of Force Development

For the acquisition of maximum torque (Tmax) and RFD data, the volunteers were seated in the chair of the isokinetic dynamometer and secured by X-shaped crossed bands on their trunk and hip. The isokinetic dynamometer was connected to a computer where the collection of electromyographic signals were synchronized with the torque curve. This was also performed through a customized physical interface developed at the Laboratory of Biomechanics and Biological Signal Processing at the School of Physical Education of the University of Brasília – UnB. Such interface is plugged to an analog-digital conversion board with a sampling frequency of 2000 Hz.

The torque data were obtained directly from the dynamometer used for calibration of the signals that were captured and stored in the computer. First, the torque signal was smoothed through a fourth order Butterworth digital filter with a cutoff frequency of 10 Hz. Then, the higher tension attempt – among the 3 attempts performed by each volunteer – was selected and converted to torque unit (Newton.meter – N.m).

To identify the RFD in moments 0-50; 50-100;100-150 and 150-200ms, the sample index closest to the value of 8.0 N.m in the torque curve of each volunteer was initially spotted. This value considers the weight of the mechanical arm and the lower limb of the volunteer, who already starts the test by producing a slight involuntary torque [20]. A few moments later, the sampling frequency (SF) of the signal was used in order to find the indexes of the digital samples that are equivalent to the points 0-50; 50-100;100-150 and 150-200ms. Considering the SF of 2000 Hz, every 50 milliseconds correspond to 100 samples. Then, the stretch corresponding to the first 50 milliseconds would be the stretch between the sample identified with a value of 8.0 N.m (x_[n]_) plus 99 samples. Proportionally we would have this reasoning for 100, 150 and 200ms, while keeping the proportions. Finally, the derivative (RFD) of the respective torque signals was calculated.

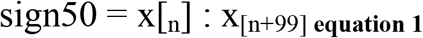

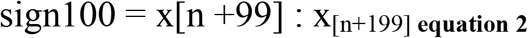

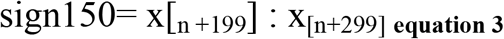

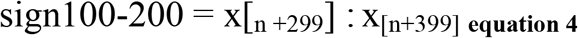

### Statistical analysis

The statistical treatment of the collected data was performed using the *IBM® SPSS software*. The evaluation of the normality of the data was made through the Shapiro-Wilk test due the number of participants to observe the normality of the values of maximum torque, RFD, MPF, RMS and AE of the electromyographic signal at the moments 0-50ms; 50-100ms; 100-150ms and 150-200ms. Next, the Spearman correlation coefficient was evaluated in order to observe the degree and significance of the correlations between the variables maximum torque (Tmax), RMS, MPF and AS and the RFD at 0-50ms; 50-100ms; 100-150ms and 150-200ms. Next, to evaluate the effect of time on the values of RFD, RMS, MPF and AE, a Friedman test for repeated measures was conducted.

## RESULTS

Table 1 presents the correlation values between the variables Tmax, RMS, MPF, AE and RFD at moments 0-50; 50-100; 100-150; 150-200ms.

**Table 1.** Correlations between the variables Tmax, RMS, MPF, AE and the RFDs at 0-50; 50-100; 100-150 and 150-200ms.

|  | RFD 0-50ms | RFD 50-100ms | RFD 100-150ms | RFD 150-200ms |
| --- | --- | --- | --- | --- |
| Tmax | 0.07 | 0.40 | 0.48 | 0.40 |
| RMS 0-50ms | 0.43 | - | - | - |
| RMS 50-100ms | - | 0.47 | - | - |
| RMS 100-150ms | - | - | 0.02 | - |
| RMS 150-200ms | - | - | - | 0.03 |
| MPF 0-50ms | **0.67\*** | - | - | - |
| MPF 50-100ms | - | 0.39 | - | - |
| MPF 100-150ms | - | - | 0.11 | - |
| MPF 150-200ms | - | - | - | 0.38 |
| AE 0-50ms | 0.42 | - | - |  |
| AE 50-100ms | - | 0.36 | - | - |
| AE 100-150ms | - | - | 0.02 | - |
| AE 150-200ms | - | - | - | 0.03 |

RFD – Rate of Force Development; Tmax – maximum torque; RMS – *root mean square*; MPF – Median power frequency; AE – Absolute Energy, \* - $p \leq 0.05$

**Figure 1.**
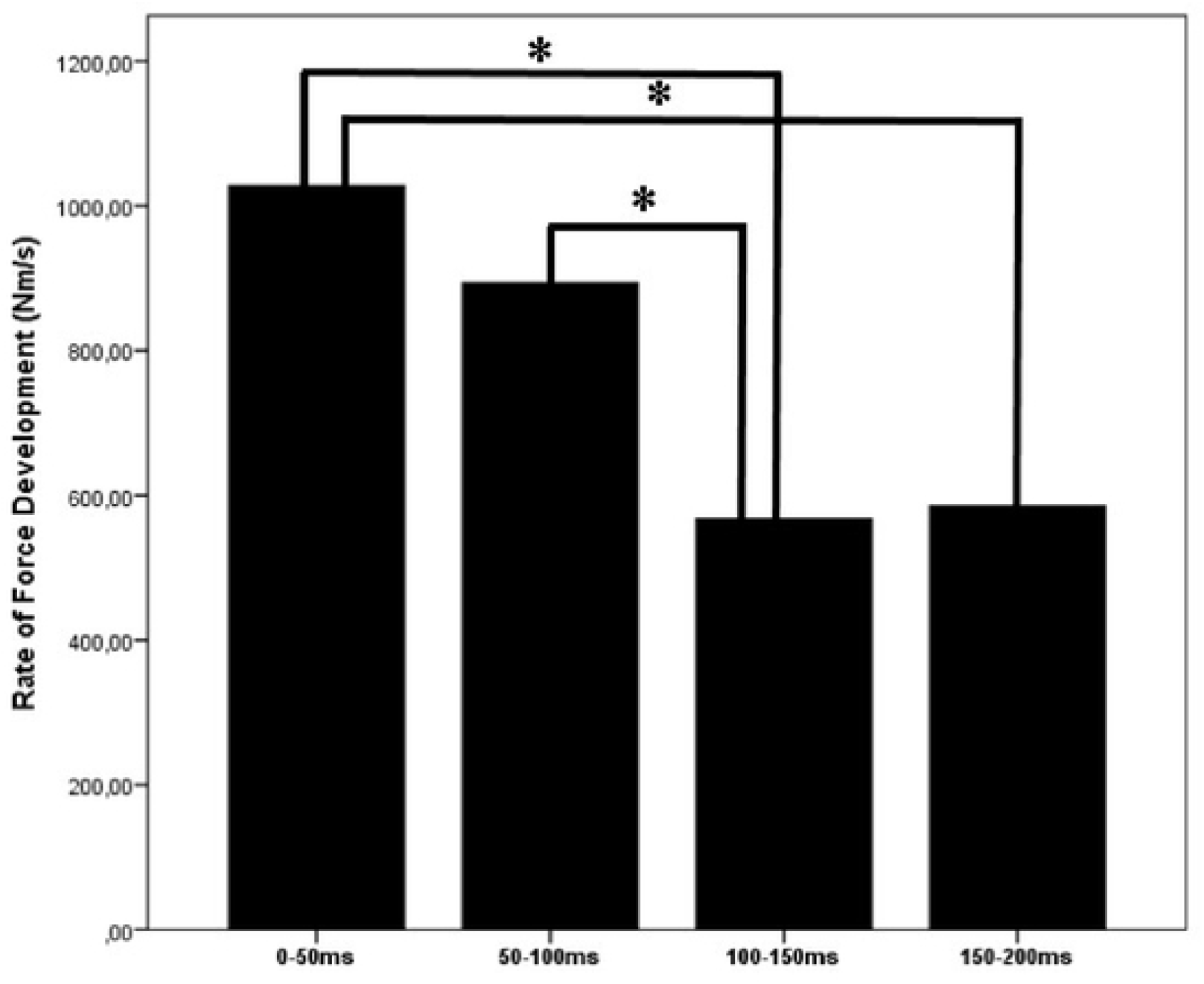
Comparison and difference between RFD values at 0-50ms; 0-50ms; 50-100ms; 100-150ms; 150-200ms. **RFD-** Rate of Force Development; **N.m/s**; **ms** – millisecond; **\*** - *p* ≤0.01 When the RFD values are evaluated, it is possible to verify a significant difference when comparing (*i*) the values of the RFD 0-50ms with the RFD 100-150ms (*p* ≤ 0.05) and (*ii*) the instant 0-50ms with the instant 150-200ms. The existence of a significant difference has also been identified when comparing the instants 50-100ms with the instant 100-150ms (*p*≤ 0.05). No significant difference was found in the comparison between RFD values at 0-50ms and 50-100ms.

**Figure 2.**
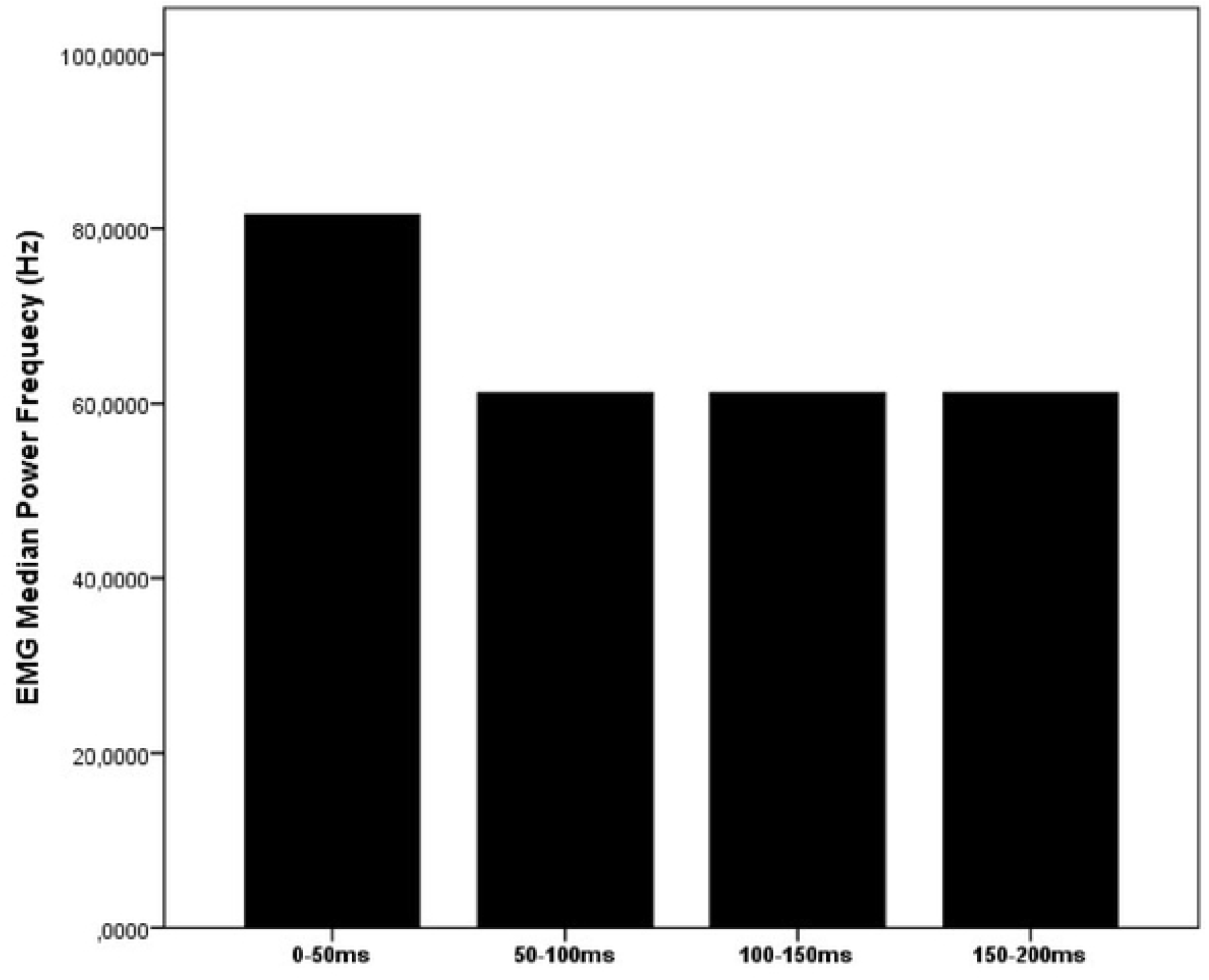
Comparison of MPF values at the instants 0-50ms; 50-100ms; 100-150ms; 150-200ms. **EMG Median Power Frequency -MPF**; **Hz-** Hertz; **ms** – millisecond. While evaluating the MPF values, it is possible to observe that there is no statistical difference between this variable at the instants 0-50ms; 50-100ms; 100-150ms; 150-200ms.

**Figure 3.**
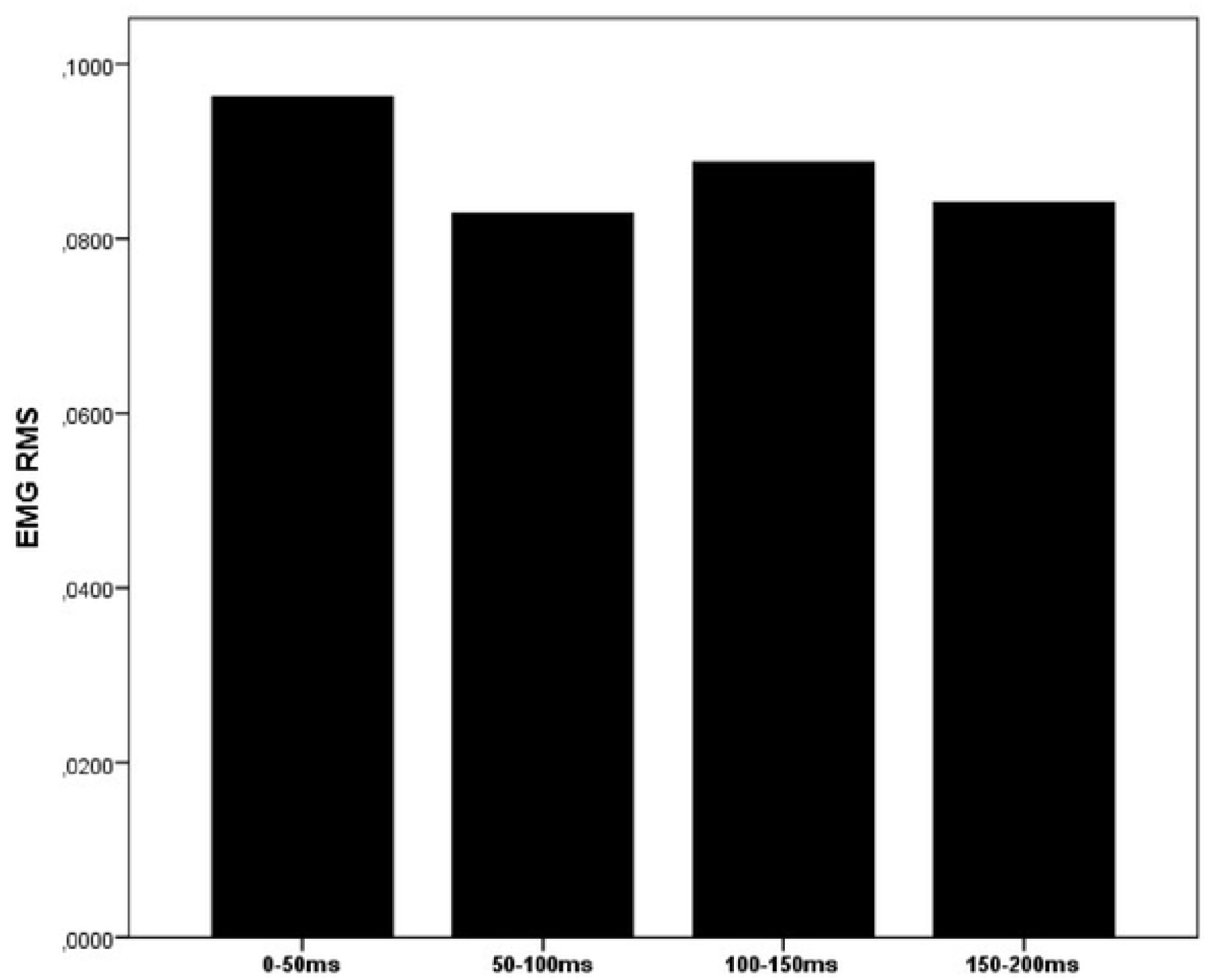
Comparison between the RMS values of the electromyographic signal at the instants 0-50ms; 50-100ms; 100-150ms; 150-200ms. **RMS –** *root mean square;* **V** – volts; **ms** – millisecond. No statistical differences were found in the comparison between the RMS values of the electromyographic signal at the instants 0-50ms; 50-100ms; 100-150ms; 150-200ms.

**Figure 4.**
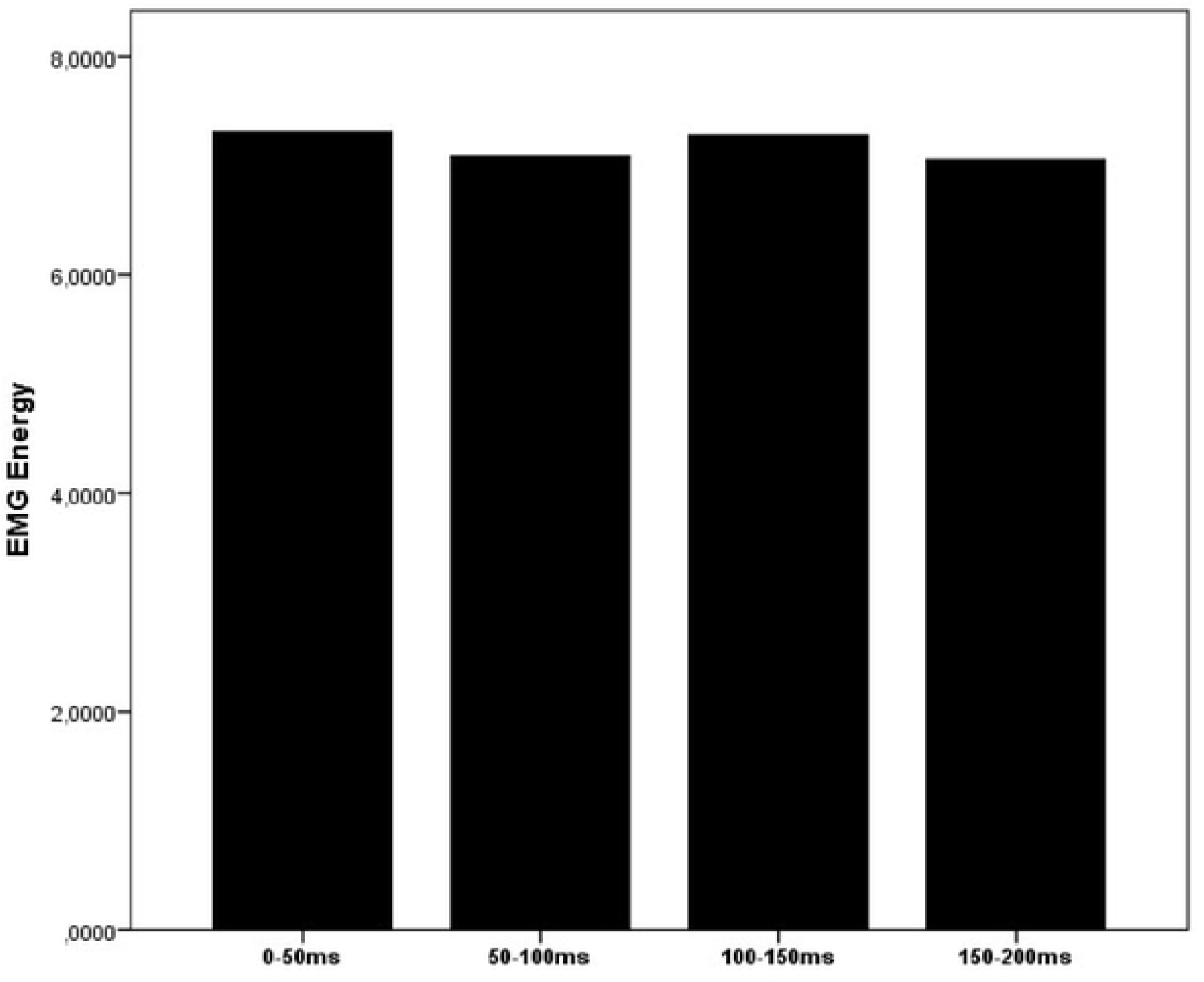
Comparison between the values of Absolute Energy of the electromyographic signal at the instants 0-50ms; 50-100ms; 100-150ms; 150-200ms. **EMG energy**; **ms** – milliseconds No significant difference was observed in the Absolute Energy of the electromyographic signal between the instants 0-50ms; 50-100ms; 100-150ms; 150-200ms.

The correlations between the values of the variables Tmax, RMS MPF and AE in the instants (0-50, 50-100, 100-150 and 150-200ms) presented in Table 1 with the values of the Strength Development Rate demonstrate a significant and strong correlation between the MPF 0-50ms and the RFD 0-50ms. For the other correlations, although they do present a degree of correlation, they do not present a significant correlation.

## Discussion

The aim of the present study was to evaluate the influence of maximal strength and neuromuscular variables – electromyographic signal energy, RMS and AE, and median power frequency (MPF) – on the rate of force development (RFD) of basketball athletes. Our main findings show there is no correlation between the maximum strength and the RFD indicator, the Rate of Force Development (RFD), during the specific moments that were studied.

Our findings also indicate that there is a significant reduction in RFD when comparing the instants 0-50ms with the instants 100-150ms and 150-200ms, as well as a significant reduction in RFD when comparing the instants 50-100ms with the instant 100-150ms.

The variation in RFD coincides with a reduction in MPF, with no statistically significant difference, at 50-100ms instants; 100-150ms; and 150-200ms, compared with the instant 0-50ms. This reduction was not observed in the values of RMS and absolute energy of the electromyographic signal, which remained at the same level at certain moments.

These results corroborate with previous studies that found no correlations of maximal isometric strength and the explosive strength indicator, RFD, in men that recreationally train for strength and wrestling [18, 22].

In our results, no correlations were observed between RMS and RFD (*r*^*2*^ = 0.43; *p<*0.05) in the same intervals. This result agrees with the previous findings which report a strong significant correlation between the RMS of the electromyographic signal and the RFD at the moments 0-50ms [12]. The study evaluated the correlation between maximal isometric strength, initial (0-100ms) and late (100-200ms) RFD and RMS of the electromyographic signal of physically active men trained for strength. The disagreements in the results suggest that it is due to the different populations evaluated, leading to the interpretation that populations more accustomed to the execution of ballistic movements, such as basketball players, tend to have determinant neural action in RFD, different from other populations.

A strong and significant correlation was observed between MPF and RFD at 0-50ms (*r*^*2*^ = 0.67; *p<*0.05), which led us to investigate it further. Previous findings corroborate the result found in our study, where it is observed that the frequency of firing of motor units presents a significant and strong correlation with the RFD of physically active people (*r*^*2*^ = 0.71; *p<*0.001). The researched authors suggest that the rate of force development is primarily governed by the frequency of firing of motor units [14].

When evaluating the behavior of RFD, MPF and RMS at the instants 0-50ms, 50-100ms; 100-150ms and 150-200ms we observed a significant reduction in RFD levels being its highest value at the instant 0-50ms. Even without presenting a statistically significant decrease in MPF (0-50ms and 50-100ms p=0.096; 0-50ms and 100-150ms p=0.31; 0-50ms and 150-200ms *p=*0.96), we observed a reduction in its levels, with its highest value observed at 0-50ms. As causes of this phenomenon, we shall mention the state of force-frequency relationship and the reduction of sensitivity to Ca^+2^ reported in previous studies [23].

The force-frequency relationship is conceptualized as the phenomenon in which the isometric force produced by the muscle increases or decreases according to the frequency of firing of the motor units. At higher firing frequencies, more Ca^+2^ is released from the sarcoplasmic reticulum resulting in increased force.

The increase occurs up to the saturation point, where even with greater release of Ca^+2^ will not result in increased strength. This is due to decreased sensitivity to Ca^+2^ in the absorption by the contractile protein troponin TNC. Previous studies mention that the greater the number of bonds the greater the force production and present the RFD values as dependent and influenced by the amount of Ca^+2^ present in the musculature capable of binding to troponin TNC [24-27].

This release of Ca^+2^ and consequently the speed of the muscle contraction response is also reduced due to a decrease in the frequency of firing which leads to a reduction in the concentration of Ca^+2^ released at each action potential [28].

It is possible to observe in our results, stability of the RMS of the electromyographic signal and a significant decrease in RFD. We understand that this phenomenon occurs due to the mechanism of not increasing the number of motor units for the maintenance of explosive force, proven by the constancy of the RMS. And yes, by the decrease in MPF. Even though there is no statistical difference between the MPF obtained in the interval 0-50ms with the other intervals studied, a fall is observed. We argue that this decrease does not need to be statistically significant to influence RFD.

In other words, while the firing frequency of the motor units decreases, within the evaluated intervals, there is also a decrease in RFD. Parallel to this occurrence, the RMS values and absolute energy of the electromyographic signal remained constant.

## Conclusion

The Rate of Force Development (RFD), is shown as a strength variable independent of the maximum strength in basketball athletes. Among the neuromuscular variables evaluated, the firing frequency of motor units (MPF) had the greatest influence on RFD. The RFD decrease coincided with the decrease in MPF and the constancy of the RMS. The results can help trainers and preparers to employ training that is focused on the variables of explosive force and frequency of firing of the motor units with training of ballistic and explosive movements for an improvement of the performance in the modality.

## Notes

### Competing Interest Statement

NO authors have competing interests Enter: The authors have declared that no competing interests exist.

